# Modelling tree growth taking into account carbon source and sink limitations

**DOI:** 10.1101/063594

**Authors:** Amaury Hayat, Andrew J. Hacket-Pain, Hans Pretzsch, Tim Tito Rademacher, Andrew D. Friend

## Abstract

Increasing CO_2_ concentrations are strongly controlled by the behaviour of undisturbed forests, which are believed to be a major current sink of atmospheric CO_2_. There are many models which predict forest responses to environmental changes but they are almost exclusively carbon source (i.e. photosynthesis) driven. Here we present a model for an individual tree that takes into account also the intrinsic limits of meristems and cellular growth rates, as well as control mechanisms within the tree that influence its diameter and height growth over time. This new framework is built on process-based understanding combined with differential equations solved by the Runge-Kutta-Fehlberg (RKF45) numerical method. It was successfully tested for stands of beech trees in two different sites representing part of a long-term forest yield experiment in Germany. This model provides new insights into tree growth and limits to tree height, and addresses limitations of previous models with respect to sink-limited growth.

**Author Summary:** Greenhouse gas emissions, in particular of CO_2_, have emerged as one of the most important global concerns, and it is therefore important to understand the behaviour of forests as they absorb and store a very large quantity of carbon. Most models treat forests as boxes with growth only driven by photosynthesis, while their actual growth depends also on many other important processes such as the maximal rate at which individual cells can grow, the influences of temperature and soil moisture on these cells, and the control that the tree has on itself through endogenous signalling pathways. Therefore, and with inspiration from process-based understanding of the biological functioning of trees, we have developed a model which takes into account these different factors. We first use this knowledge and additional basic assumptions to derive a system of several equations which, when solved, enable us to predict the height and the radius of an individual tree at a given time, provided that we have enough information about its initial state and its surroundings. We use the Runge-Kutta-Fehlberg mathematical method to obtain a numerical solution and thus predict the development of the height and radius of an individual tree over time under specified conditions.

## 1 Introduction

Forests are an important component of the global carbon cycle and are currently a major sink of atmospheric CO_2_ ([1]; [2]). Being able to predict the future response of forests is therefore of great interest. Many models have been used to address this issue, but they are almost exclusively carbon source-driven, with plants at any particular location treated as a box, or boxes, of carbon mainly driven by photosynthesis (e.g. [3]; [4]; [5]). However, it is likely that many other factors, such as the intrinsic limits of meristems and cellular growth rates, as well as control mechanisms within the tree, have large influences on forest responses ([6]; [7]). A few research groups have addressed the issue of sink-limited growth in a modelling context. The potential for sink-limited growth to affect carbon storage and treeline position was addressed by [8] using a global vegetation model. However, their approach was highly empirical and only addressed temperature limitations. A more mechanistic approach was presented by [9], in which a tree-level carbon balance model was constructed with both carbon source and sink parameterisations. The sink parameterisations were based on thermal-time and included different ontogenetic effects between tissue types and xylogenetic processes for secondary growth. However, while this paper makes a significant contribution, the various parameterisations were very simply incorporated, with no effect of moisture on wood growth, fixed durations for xylem enlargement, and no overall tree growth across years. [10] also presented a tree-level model parameterisation that addressed the effect of growth processes independently of photosynthesis, and in this case looked particularly at soil moisture effects. However, they did not explicitly treat meristem growth, but instead used modified allocation coefficients depending on temperature and soil water. Here we present a new framework for addressing tree growth responses to environmental change, building on knowledge of tree physiology to develop approaches for predicting the development of an individual tree, and thereby enabling a better understanding of forest responses to environmental change than purely source-driven models can achieve, as well as addressing the limitations of previous sink-limited approaches.

In this paper we suggest that this objective can be realised using differential models, that is to say evolution models using differential equations. We propose here a first differential model taking into account control mechanisms and the intrinsic limits of meristems and cellular growth within trees together with the carbon balance. The paper is presented as follows. First we make several hypotheses concerning physiological considerations, then we give a description of the model. We show how the model is used to predict tree height and stem volume in two sets of three different stands of beech trees. Finally, we discuss the results and the behaviour of the model in several situations.

## 2 Model and framework

### 2.1 General Considerations

To describe the development of the tree, we suppose that it has two types of meristems, apical and lateral, and that the apical meristem increases the height through sustaining primary growth, while the lateral meristem (i.e. the vascular cambium) increases the radius through sustaining secondary growth. We recognise that trees will usually have many separate apical meristems distributed across many branches, but for convenience we treat these together as one apical meristem; we also ignore apical root growth for now. We further assume that the stem can be represented by a cylinder and that the crown (i.e. the branches and foliage) occupies a cylindrical volume and has dimensions that are proportional to the stem dimensions, i.e. the crown depth is proportional to the height of the tree stem and its radius to the radius of the tree stem. In this way we need only obtain information about the radius *r* and the height *h* of the stem to describe the growth of the tree. Future developments will include introducing other crown and stem shapes, as well as root meristems. Here our objective is to keep the model as simple as possible in order to explore the realism of its fundamental assumptions.

In order to derive the dynamics of *r* and *h* we need three types of equations:

- Constitutive equations - related to the structure of the tree;
- Carbon balance equations - related to the balance of carbon between the tree and its environment, and the balance within the tree; and
- Control equations - representation(s) of the intrinsic controls that the tree has on itself.

### 2.2 Constitutive equations: allometric relationships

Physiologically, plant growth is sustained by meristems producing new cells which subsequently enlarge and increase in mass ([11]). We suppose that for each type of meristem, growth is proportional to its volume, that is to say that the mass of carbon allocated to a meristematic region will depend proportionally on the volume of meristem, which determines the rate of new cell production. We also suppose that a tree controls the activities of the meristems and therefore the relative demand for carbon *between* the meristematic regions: it can favour either height or diameter growth in this way, depending on environmental signals. Moreover, we suppose that this is the only control the tree has on its growth. With this hypothesis we soon get:

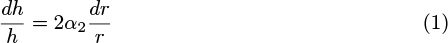

where *α*_2_ is the ratio of activities between the apical and lateral meristems, i.e. when the activity of the lateral meristem produces 1 *g* of carbon growth, *α*_2_ *g* of carbon growth occurs from the apical meristem. It is the control parameter.

Note that we suppose here that the volume of the apical meristem is proportional to the top surface area of the (cylindrical) stem and that the volume of the lateral meristem is proportional to the lateral surface area of the stem. This hypothesis is used as it is more realistic than assuming that the volume of the lateral and apical meristems are proportional to the volume of the tree, i.e. that the thickness of the lateral meristem area increases proportionally with the radius and the thickness of apical meristem increases proportionally with height. Although even under this alternative hypothesis we would get the same type of equation with *α*_2_ being the proportion of growth per unit of volume of apical meristem relative to proportional lateral growth, only adding a numerical coefficient in front of *α*_2_.

Knowing the definition of the volume and integrating we get the following constitutive equations:

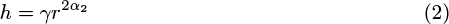

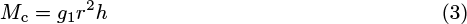

We recognize Equation (2) as the usual allometric relationship where *γ* is a constant. As it is true for any time *t*, it can be expressed as 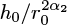, with *h*_0_ and *r*_0_ the height and radius at some time origin *t* = 0.

Equation (3) is simply the definition of the volume of the stem plus canopy (containing the branches and leaves) in relation to its mass, *M*_c_. *g*_1_ is the proportionality ratio that takes into account the density of the stem, the density of the branches and foliage, their contribution to the mass of the tree, and a geometric factor such that *g*_1_*πr*^2^*h* is the mass of the tree. *g*_1_ can therefore be found using the proportionality coefficients we assumed between crown radius and stem radius and crown height and stem height, and their relative densities.

### 2.3 Control mechanism of the tree

We want now to take into account the intrinsic control that the tree has on itself, parameterised in our model by *α*_2_, the ratio between the activities of the apical and lateral meristems. For this we need to consider what happens physiologically (for convenience we use teleological terminology): the tree uses internal controls to be able to adapt its shape and physiology to the surrounding environment, for instance other competing trees (e.g. [12]), through phytohormonal signalling networks ([13]; [14]). Many behaviours could be taken into account, but here we focus on the behaviour of a tree competing for light by trying to grow taller faster than its competitors by increasing *α*_2_. The modelled tree is assumed to detect the presence of surrounding trees by sensing the ratio of downwelling red radiation (i.e. wavelengths between 655 and 665 nm) to downwelling far-red radiation (i.e. wavelengths between 725 and 735 nm). A low ratio signals potentially more neighbouring trees and therefore a potential threat of shading. In that case the tree reacts by increasing the activity of the apical meristem relative to the lateral meristem, while a higher ratio potentially means no threat and so the tree reacts by balancing the activities of the two meristems as it needs to compete less for light ([15]). We hypothesise that this reaction can be modelled as:

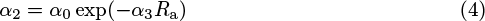

Where *α*_0_ is the highest limit for *α*_2_ beyond which the tree breaks due to mechanical failure, and *R*_a_ is the ratio of received downwelling red:far-red ratio. *α*_3_ is a scaling parameter such that *α*_0_ exp(−*α*_3_) is the value of *α*_2_ when the tree is unshaded (i.e. when it detects no other tree around).

This approach is based on the findings of several studies (e.g. [15]; [16]). In particular, [16] observed the laboratory behaviour of two similar young trees grown for 21 days with the same intensity of photosynthetically active radiation (PAR), but different red:far-red ratios. They experimentally obtained a relationship which, extended by limited development for short trees, is coherent with the relationship we give in Equation (4).

A potential limitation to our approach is that we may not have access to direct measurements of *R*_a_. For a lone tree this would not be a real problem: the allometric relationship would be constant in our model and the red:far-red ratio maximal. For individual trees growing in a stand in a forest, however, the situation is different as the trees shade each other. In order to overcome this problem, we can try to obtain *R*_a_ from more accessible variables. Intuitively, the ratio should depend on the density of trees and the area they shade horizontally. This ratio should also allow a finite maximum when the density tends to 0, that is to say there are no neighbouring trees. Therefore, we could propose the following dependency:

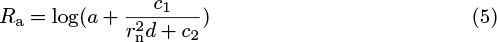

With *d* the density of trees (ha^−1^), *r*_n_ the average radius of the neighbouring trees (m), and *a*, *c*_1_, and *c*_2_ are three constants such that 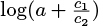 is the maximal value of the ratio (i.e. when there are no trees around). This formula is inspired by several studies (e.g. [12]; [17]), where red:far-red ratios were measured together with the density of the stands, and which suggest that for young trees the ratio has a logarithmic increase with the inverse of the density. However, it is likely that this is our most limiting relationship as we are here trying to model the internal control that the tree has on itself, which is probably the most complicated phenomenon to take into account, and likely the least-well understood component of our model (cf. [14]; [18]). It is also easy to see that the *r*_*n*_ variable makes this formulation challenging to use: even with an average value, finding the solution of 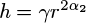 becomes difficult and in practice would increase the complexity of the differential equations. Also, it should be taken into account that this formula becomes very sensitive to any small variation of *r* due to measurement error or the consequences of a particular year’s conditions. This is the reason why, to keep the model simple, we consider that when we do not have access to measurements of the red:far-red ratio, *α*_2_ is constant for each stand, as it would be for a lone tree. When the value of this constant will be needed, it will be estimated as a typical value for the species we consider from a stand with similar density.

### 2.4 Carbon Balance

The variation of carbon in the tree over time can be written as the sum of the carbon used for volume growth and the carbon stored:

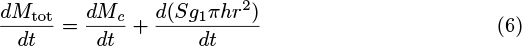

where *M*_*c*_ is the carbon used for volume growth, *S*_*c*_ = *S*_*g*_1__*πhr*^2^ the carbon stored, and *S* is defined as carbon stored per unit volume of stem. This distinction has a physical interpretation and therefore in this model the density of the tree can then be seen as a standard density related to the carbon used for growth plus an increase of density due to the storage.

We also know that the variation of carbon is given by:

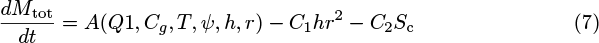

where *A* is the carbon assimilation due to gross photosynthesis minus a foliage respiration part proportional to photosynthesis, while *C* is a constant grouping a part of the respiration and a litter component proportional to the volume, together with the geometrical factor *g*_1_*π* (such that *C*_1_/*g*_1_*π* is the respiration and litter component constant). We assume here that there is no other litter component, that is to say that litter is completely proportional to the volume of the stem. We note that this also means that the litter is proportional to the volume of the crown with our previous assumptions. Finally, *C*_2_ is the same type of constant but for the storage. *A* is assumed to depend on PAR (Q1), the concentration of CO_2_ in the atmosphere (*C*_g_), temperature (*T*), water potential (*ψ*), and the dimensions of the tree (*h* and *r*).

Therefore:

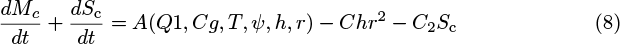

The system is so far incomplete as there needs to be a relationship that controls which part of the carbon is allocated to growth and which part is allocated to storage Also, the volume growth of the tree is sustained by meristematic cell division and enlargement, with a maximal rate determined by intrinsic physiological limits or environmental controls.

We assume that the tree grows in volume as much as it can, as long as it keeps enough storage to avoid being in danger, such as for repairing damage and surviving poor growing seasons. Therefore, if there is enough carbon assimilation then the rate of growth is equal to the meristem-sustained growth-rate limit and the storage can increase using the difference of these two. Also, if the carbon assimilation rate becomes too low but there is enough storage, then the storage delivers carbon to the meristematic region to maintain a rate of growth equal to its maximal sustainable rate. Then density decreases as the carbon of the storage is used to maintain the volume growth. This is likely to occur for instance when buds appear while photosynthesis is not yet high enough to support the maximal growth potential due to meristem reactivation (note that bud-burst and other seasonal phenological phenomena are not yet treated by the model). However, if there is not enough carbon assimilation or storage then the rate of growth is equal to the carbon assimilation rate and the storage per unit volume remains constant.

This behaviour can be translated into evolution equations. As growth occurs at a cellular level we can assume that the maximal meristem growth is proportional to the volume of meristems. We denote this maximal meristem-sustained growth per volume by *R*_max_, the volume of the meristems by *V*_me_, and the storage limit under which the tree is considered in danger by *S*_1_.

If there is enough light or storage, i.e.
if 
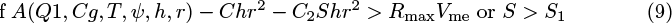

then:

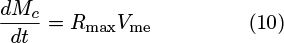

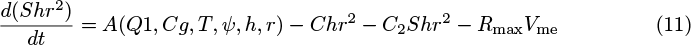

but if 
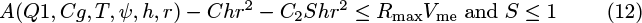

then

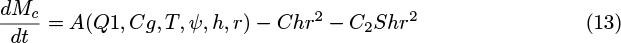

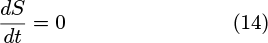

### 2.5 Modelling Photosynthesis

We assume the following dependencies of photosynthesis on water potential, atmospheric CO_2_ concentration, PAR, temperature, radius, and height:

1. Photosynthesis increases with CO_2_ concentration and PAR exponentially and tends to saturate:

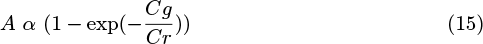

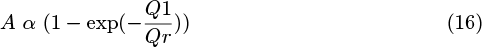
2. As a first approximation photosynthesis of each leaf decreases linearly with a reduction in water potential. We suppose also that it decreases linearly with height:

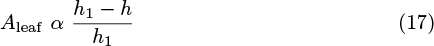 Where:

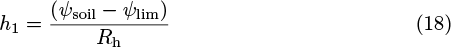 With *ψ*_soil_ the water potential in the soil and *ψ*_lim_ the limit water potential below which any photosynthesis cannot be performed due to a lack of turgor pressure in the leaves and increased probability of (xylem) cavitation. *R*_h_ is the hydraulic resistance per unit length. So, integrating across the full tree, we get:

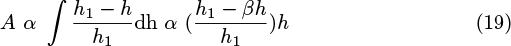 Note that *β* is a numerical coefficient that only depends on the ratio between the crown depth and the height. If we suppose that the crown starts at around half the height of the tree then *β* = 3/4. Also, the dependency of both height and water potential is given above.
3. We assume a dependency with temperature given by an asymmetric parabola:

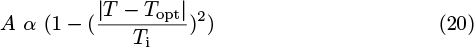

With *T*_i_ being 21°C if *T* > *T*_opt_ and 25°C otherwise. We set here *T*_opt_ = 19°C. These values are inspired by previous studies (e.g. [19]).
4. We suppose that photosynthesis is proportional to the surface area of the crown, and so proportional to *r*^2^. Overall we get:

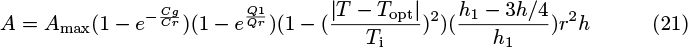

### 2.6 Maximal meristem-sustained growth

Intuitively, as meristem-sustained growth occurs at a cellular level, we assume that the limit to meristem-sustained growth per volume increases with temperature up to a certain temperature and increases with water potential (cf. [20]). Therefore, we assume the following dependencies:

1. The limit to meristem-sustained growth per volume increases with temperature using a standard *Q*_10_ formulation: 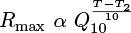
2. The meristem-sustained growth limit decreases with height as: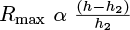

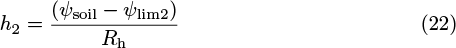 With *ψ*_soil_ the water potential in the soil and *ψ*_lim2_ the limit water potential below which meristem-sustained growth cannot occur. Physically this gives us a maximal height *h*_2_ due to the limit on meristematic growth. We define therefore *R*_max0_

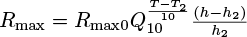
3. As stated previously, we suppose that the volume of meristems is expressed as: 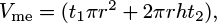 where *t*_1_ is the effective thickness of the apical meristem and *t*_2_ is the effective thickness of the lateral meristem.

### 2.7 Final evolution equations

Using the previous elements we finally get:

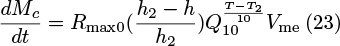

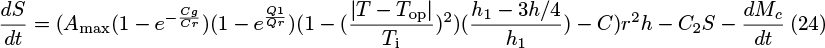

or

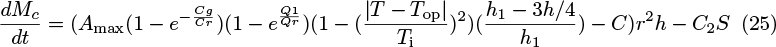

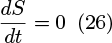

depending on whether carbon supply from photosynthesis plus storage is sufficient to meet demand for growth (Eqns. 23 and 24) or not (Eqns. 25 and 26).

### 2.8 Deriving the values of the parameters

So far we need to know the values of the following physiological parameters to use the model:

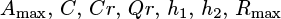

The most accessible are probably *Cr* and *Qr* These are usually well known and can be obtained by looking at the curve of photosynthesis activity as a function of atmospheric CO_2_ mixing ratio (in ppmv) or as a function of PAR, respectively.

*A*_max_ can be obtained by direct measurements of the carbon exchange at a given height, ambient CO_2_ concentration, temperature, PAR, and compensating for respiration as in [21]. Acting similarly and measuring litter with litter traps gives access to *C*. These values can also be deduced from net primary production and gross primary production and the typical litter flux rates. Note that here we use data coming from forest stands, although in the model we suppose that we have an individual tree without shading because data for lone standing trees are seldom available. This means that we may underestimate the growth of isolated trees, but at the same time the values of *A*_max_ and *C* obtained enable to take into account the shading effect by neighbours of a forest with the same density, which is a hidden parameter of this model (i.e. not explicitly taken into account). Also, depending on the typical time length of the study, we choose a model timestep and therefore we use averaged values of the parameters on this timestep. In this analysis, we chose one year as the timestep although smaller timesteps could be considered.

Finding *h*_1_ is equivalent to finding the minimal water potential that can sustain photosynthesis. *h*_2_ is its equivalent with respect to growth. Combined, these two parameters take into account all water potential-dependant limits in trees such as cavitation in the xylem conduits, minimal turgor potential to maintain the shape of the leaf and enable stomatal opening, etc. While *h*_1_ (or equivalently *ψ*_max1_) has been extensively studied (e.g. [22]; [23]; [24]), and is believed to be equal around 90 m for redwood trees ([23]), *h*_2_ seems to be less known. However, knowing *h*_1_ and the other parameters as well as the effective limit height for the considered species, we can deduce *h*_2_ relatively easily.

*Q*_10_ is a physiological parameter that can be found by studying the activity of meristem cells in a controlled environment. We can find *R*_*max*0_, at least approximately, by looking at the tree rings of many trees across several geographical locations and assuming that the largest ones correspond to the maximal increment *dr* achievable in a year, which is directly linked to *R*_max0_, the maximal meristem-sustained growth rate under perfect temperatures and water potentials. Then we can deduce the value of *R*_max0_, or at least its order of magnitude.

**Table 1.** Numeric values of the parameters used in the model.

| *Symbol* | *Value* | *Quantity* |
| --- | --- | --- |
| $g_1$ | $477 \text{ kg.m}^{-3}$ | Density of the equivalent cylinder modelling the tree |
| $C/(g_1\pi)$ | $72 \text{ kg.m}^{-3}.\text{y}^{-1}$ | Respiration and litter coefficient |
| $C_2/(g_1\pi)$ | $72 \text{ kg.m}^{-3}.\text{y}^{-1}$ | Respiration coefficient for the storage |
| $t_2 R_{\max 0}$ | $0.0201 \text{ m.y}^{-1}$<br>corresponds to a maximal radius expansion of $1.5 \text{ cm.y}^{-1}$ | Reference maximal meristem-sustained lateral growth per area independent of water potential and temperature |
| $t_2/t_1$ | 1 | Ratio between lateral and apical meristem thickness |
| $\beta h_1$ | $67.5 \text{ m}$ | Limit height of occurrence for photosynthesis |
| $h_2$ | $47 \text{ m}$ | Limit height of occurrence for meristem-sustained growth |
| $S_1$ | 0.2 | Critical value of the storage per unit volume |
| $A_{\max}/g_1\pi$ | $206 \text{ kg.m}^{-3}.\text{y}^{-1}$ | Maximal assimilation of carbon per unit volume by photosynthesis |
| $Q_r$ | $1000 \text{ W.m}^{-2}$ | Characteristic PAR dependency |
| $\alpha_2$ | 0.34-0.43 | Ratio of meristematic activity |
| $C_r$ | 500 ppm | Characteristic CO<sub>2</sub> mixing ratio dependency |
| $\beta$ | 3/4 | Geometrical integrand for $h_1$ |
| $T_{\text{op}}$ | 18 °C | Optimal temperature for carbon assimilation |

## 3 Results

The model described above is ordinary, first order, non-linear, and has no widely known explicit analytical solution in the case where 2*α*_2_ is not an integer. Therefore, it was solved numerically by the Runge-Kutta-Fehlberg (RK45) method ([25]). Coherence both with the analytical solution in simplified cases and with other solving methods (i.e. the trapezoidal rule and numerical differentiation formula) was tested.

As some parameters can depend, *a priori*, on the considered species, we derived the parameters explained previously for beech trees from the data provided in [21] and [26]. Beech trees were chosen because of their importance in European forests, especially in Germany where they represent more than 15% of the forest and in France where they comprises 15% of the non-conifer forest. Therefore, a great number of studies have focussed on beech trees, enabling easier comparisons with our current study.

As the computation of the parameters is based on independent measurements that have sometimes a non-negligible margin of error, we allowed the parameters thus obtained to vary with a 20% margin to account for this error, and to allow a small adaptation of the model (as it is only a simplification of reality) by fitting them with reference measurements of a beech tree stand in Fabrikschleichach, Germany. As *R*_max0_ was more coarsely estimated we allowed it a variation of 50% in order to maintain the right order of magnitude. Also, as no precise data on the soil water potentials over the tested period A.D. 1870 to 2010 were available for these stands, we allowed a 50% variation of *h*_2_, the height where meristem-sustained growth cannot occur due to cell turgor limits. We obtained then a final vector of parameters for our model using a non-linear least-square method (trust-region-reflective algorithm: http://uk.mathworks.com/help/optim/ug/equation-solving-algorithms.html).

We then used our model to predict the time evolution of height and volume of an average tree in several stands of beech trees. It should be emphasised that there is a substantial difference between fitting and prediction. Fitting a model to some data gives a partial understanding of the data but does not usually enable prediction for another tree or even for future data points as the fitting is done on a restricted set of measurements. Prediction, on the other hand, is much harder to achieve as it supposes the use of previously-derived parameters and environmental information (e.g. air temperature, soil moisture, CO_2_ concentration, etc.) to predict the measurements. Usually most models are fitted rather than predicting as it is much harder to predict anything without any fitting, although when it works it gives much more information: a better understanding and a reliable way to deduce future measurements before they occur even for other trees (e.g. height or volume prediction).

We first considered three stands of beech trees that are part of a long-term experiment on forest yield science performed by the Chair of Forest Growth and Yield Science at the Technische Universität München. The stands are in southern Germany, about 60 km from Würzburg in the heart of the Steigerwald, a richly forested hillside, near a small village named Fabrikschleichach (49.9 °N; 10.5 °E). They cover roughly 0. 37 ha, are located within immediate proximity of each other (i.e they have experienced the same environmental conditions), and differ only in silvicultural management, especially thinning with consequences for the development of stem densities during the experiment. The beech trees were planted and then measured at irregular intervals averaging 8±3 (mean±standard deviation) years from an age of 48 yr in A.D. 1870 to 168 yr in A.D. 1990. For each stand we considered the averaged height of the 100 trees with the largest stem diameter at breast height. The first stand was the reference stand used to partially fit the parameters (within the 20% margin).

As expected, our partially fitted model agrees well with the measurements from this stand (Fig. 1 a; *R*^2^ = 0.975). Predictions with the model were then performed for the two other stands (Fig. 1 b,c).

**Figure 1.**
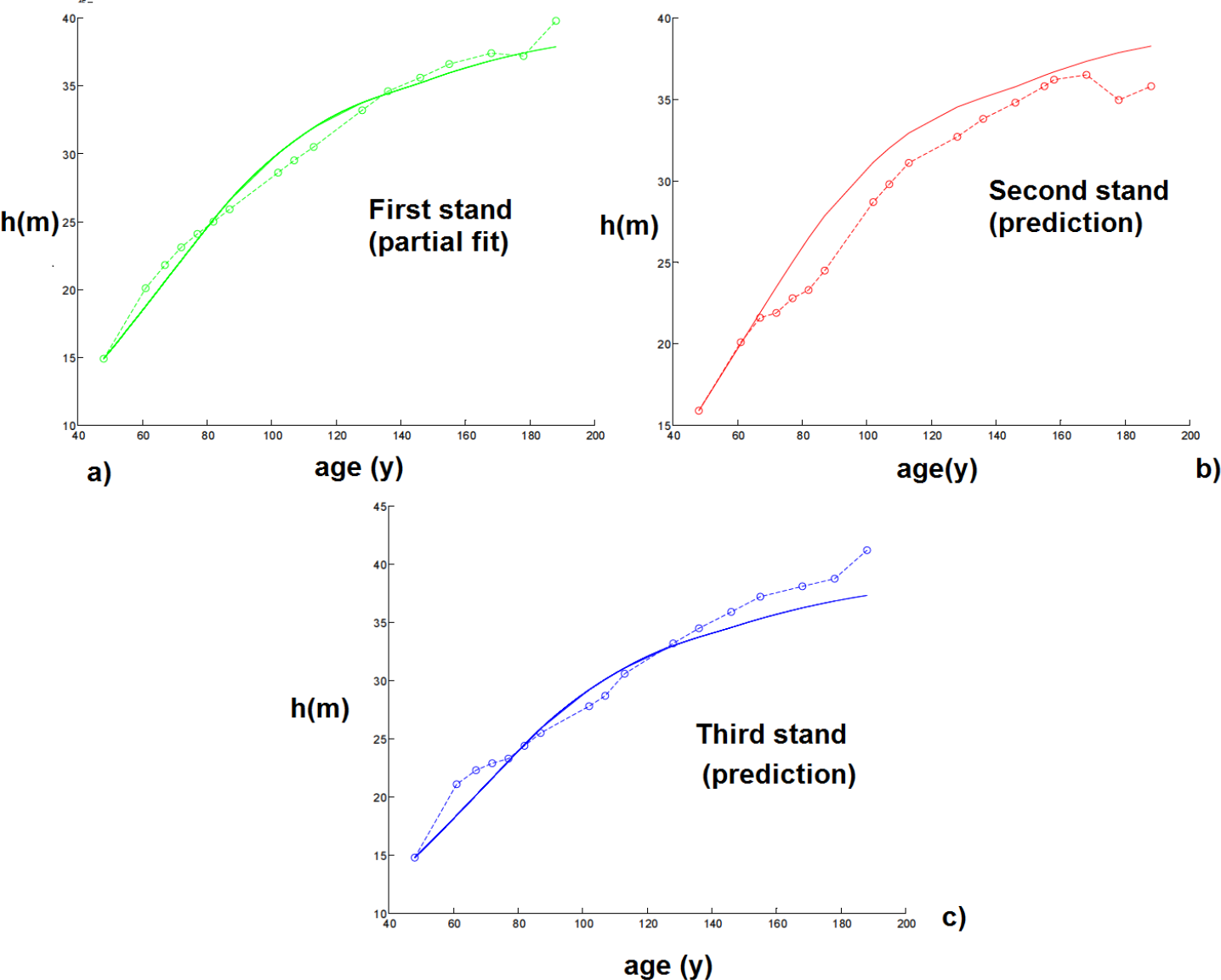
Size of the modelled tree with age at the first site. Continuous curves are computed with the model while circles correspond to experimental measurements. a) represents the results of the partial fit on the reference stand. b) and c) represent the results of the prediction with the parameters obtained by partial fitting.

The results give a very good prediction for the third stand (*R^2^* > 0.94), and a small, although non-negligible, error for the third one (*R^2^* = 0. 905). However, this could be at least partially explained by the change in the density of trees during the 120 yr due to different rates of thinning in the different stands (i.e. from 3,500 tree *ha^−1^* to 200 tree *ha^−1^* in the first stand, 6,400 tree *ha^−1^* to 200 tree *ha^−1^* in the second, and 2,400 tree *ha^−1^,* to 200 tree *ha^−1^* in the third). Therefore, the red:far-red radiation ratio changes with time in the real forest while it was assumed constant in our model.

To test the generality of the model, it was also used to predict the height of dominant trees in three other stands at a different site in the Spessart, Hain (50.0 °N; 9.3 °E), a low wooded mountain range in central Germany, located approximately 100 km from the first site (Fig. 2). As data were missing concerning the water potential during the growth period from A.D. 1870 to compare with the first stand, we allowed the model to adapt its value of *h_2_* within 20% by keeping the parameters derived previously and fitting only this one on the first of the three new stands. Then the set of parameters were used to predict the heights of the three stands according to the model. Again, to avoid the potential error induced by the different variations of the red:far-red ratios within the stands, we restrict our analysis to data when the density was lower than 1,000 tree *ha^−1^.* As before, the predictions are in strong accordance with the observations for all three stands, with a nearly exact correspondence with the measurements (*R*^2^ = 0.98, *R*^2^ = 0.91, *R*^2^ = 0.97 respectively), except for the second stand.

We should note that the correspondence is very close but not exact, and that the data we have might limit the accuracy: firstly, from our equations we can see that there is a propagation of error from the initial conditions. An error on the initial height of 1 m could imply an error of the predicted height of up to 2.5 m after 50 years even if the model were perfect. Also, no data were available for the difference of water potentials between the stands on a same site, while it seems logical that for a certain density of trees, the stands with higher density will have less water available per tree. Therefore, this could induce a small error in *h*_2_ and *h*_1_ that could be corrected if we knew the average water potential during the growth period in each stand.

Finally, mortality is not addressed by the model, although the model does produce cessation of growth under stressful circumstances. Therefore tree mortality due to critical events (e.g. disturbance, pathogens, etc.) other than a limit on growth due to the external conditions, can create discontinuities in the data which are not captured by the model. Although the continual use of the 100 trees with largest DBHs tends to average this discontinuity (as the probability that a large change in those 100 trees occurs in one year is low), this might have created another limit to the accuracy. For instance, we note that we have observed in the 2010 measurements, i.e. after 188 years, the death of several of the 100 trees with the largest DBHs, which seems to have caused a decline of the average height of the 100 living trees with the largest DBHs as taller trees were replaced by smaller trees in this group, which is a result of size-related mortality dynamics (cf. [27]).

If we wanted to address further this question, especially if we wanted to simulate many individual trees that interact together in a forest, we could multiply the height *h* by a random process *M*(*t*) that would be equal to 1 when the tree is alive and 0 when the tree is dead. Then the probability p(M(t)=0|M(s<t)=1) of transition between a living tree and a dead tree could be derived using experimental measurements and even be a function of the size of the tree. So far, though, this doesn’t seem to be needed in the present application.

So far, the only things we assumed to be known for predicting the height are the external variables (i.e. temperature, water potential, PAR, and CO_2_ concentration), the species of the tree, and the allometric relationships for each stand. No other knowledge about the trees was used to perform the predictions. Measurements in themselves were used only to compare our predictions with reality.

**Figure 1.**
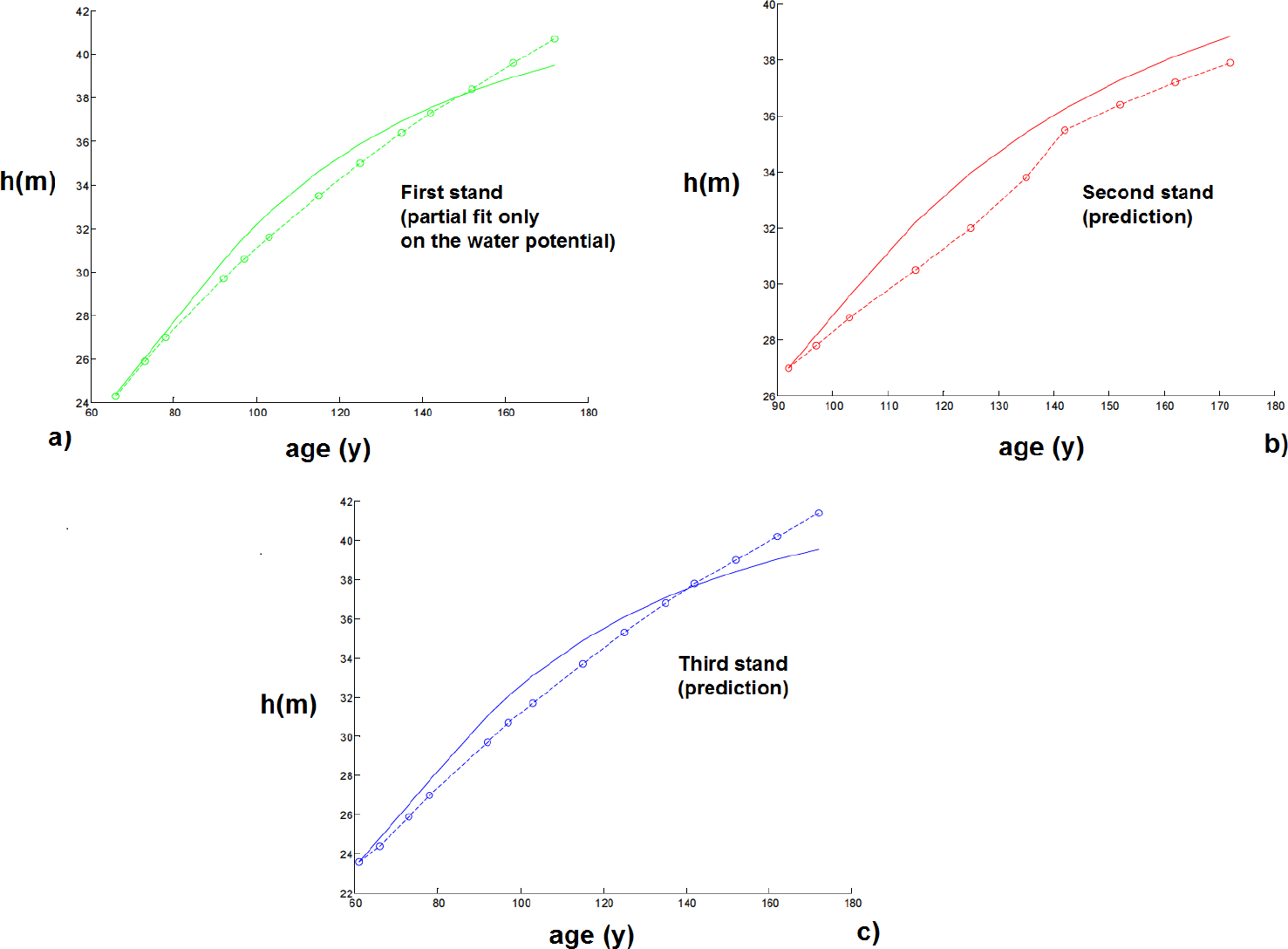
Plot of the size of the tree with age at the second site. Continuous curves are computed with the model and start when the tree density is lower than 1,000 tree *ha*^−1^, while circles correspond to experimental measurements. Parameters for beech trees are obtained from the first site and *h_2_* was allowed to vary up to 20% to compensate for the lack of data concerning water potential from A.D. 1870 on the two sites. a) to c) represent respectively the results for stands 1 to 3.

We could avoid assuming known allometric relationships for each stand and try to estimate them as described in Section 2.3 (“Control mechanisms of the tree”). In that case we would still get good results with *R*^2^ around 0.8 – 0.9. The small difference of accuracy between this case and the previous one is mainly due to the model becoming more sensitive to measurement error when we estimate the allometric relationship, and also to the estimated allometric relationship being a constant as explained in Section 2.3, whereas the allometric relationship in reality changes with changes in tree density and therefore time. Lastly, our model for *α*_2_ is probably too simple as we also explained in Section 2.3. Nevertheless, we can note that the accuracy of the results is still very good considering that we try here to predict the heights of three stands without knowing anything but the species, the CO_2_ concentration, the temperature, the soil moisture, the PAR, the number of trees per ha, and the initial tree dimensions. This last point also underscores an additional motivation to build such a relatively simple model: collecting data about the external variables can be difficult. Therefore, data can be limiting and in order to be practical the model should stay as simple as it can relative to the required data, provided that it can still give accurate results.

We chose to predict height as there is usually a relatively small error of measurement in height compared to other variables. It should be noted that the same procedure is possible taking instead the stem volume as the quantity to predict. However, the results would be less precise because they will be susceptible to cumulative measurement errors of the radius and the stem height, among other reasons, but they would still be accurate (the highest error is around 25% after 75 years) and are presented in Fig. 3. Also, we chose to predict height instead of radius or diameter as the measurements performed on the radius were the DBH which might be different from the equivalent cylindrical diameter, especially for the 100 trees with the largest DBH, and this would have added an artificial error of measurement.

### 3.1 Notable Behaviour

The results of this model are very encouraging and are not limited to predicting the height of dominant trees but show notable qualitative behaviours in other respects.

Firstly, it is known that forest stand growth dynamics have significantly changed since 1870, as shown in [28], and many experiments have been conducted to measure the impact of climate and CO_2_ change on forests. Climate and CO_2_ change also shows an impact on this individual tree model, as the model depends on external conditions such as temperature and soil moisture, as well as CO_2_ concentration. For instance, using the model with a constant temperature CO_2_ concentration (both from 1870) rather than the actual temperatures CO_2_ concentrations gives biased results and a slower growth of trees (Fig 4). This seems coherent with both the idea that climate change had an impact on forest stand growth and that present forest stands are growing more rapidly than comparable stands before.

Also, if we consider a small tree in a forest, shaded by the canopy above, it would receive only a small amount of light. Therefore the model would predict a very limited maximum height and the tree would remain near this height with very small growth until a big tree nearby falls. Then the light received would be higher and enough for the tree to grow taller very quickly. If the tree were very close to the dead big tree then the aperture is large enough and the tree would reach the canopy. If not, and if the tree is still significantly partially shaded compared to the rest of the trees, then its limit height would be lower than the forest height and it would reduce its growth progressively until it were close to this height. This behaviour would seem to be qualitatively in accordance with reality [29], but due to a lack of data we have not been able test this quantitatively.

**Figure 3.**
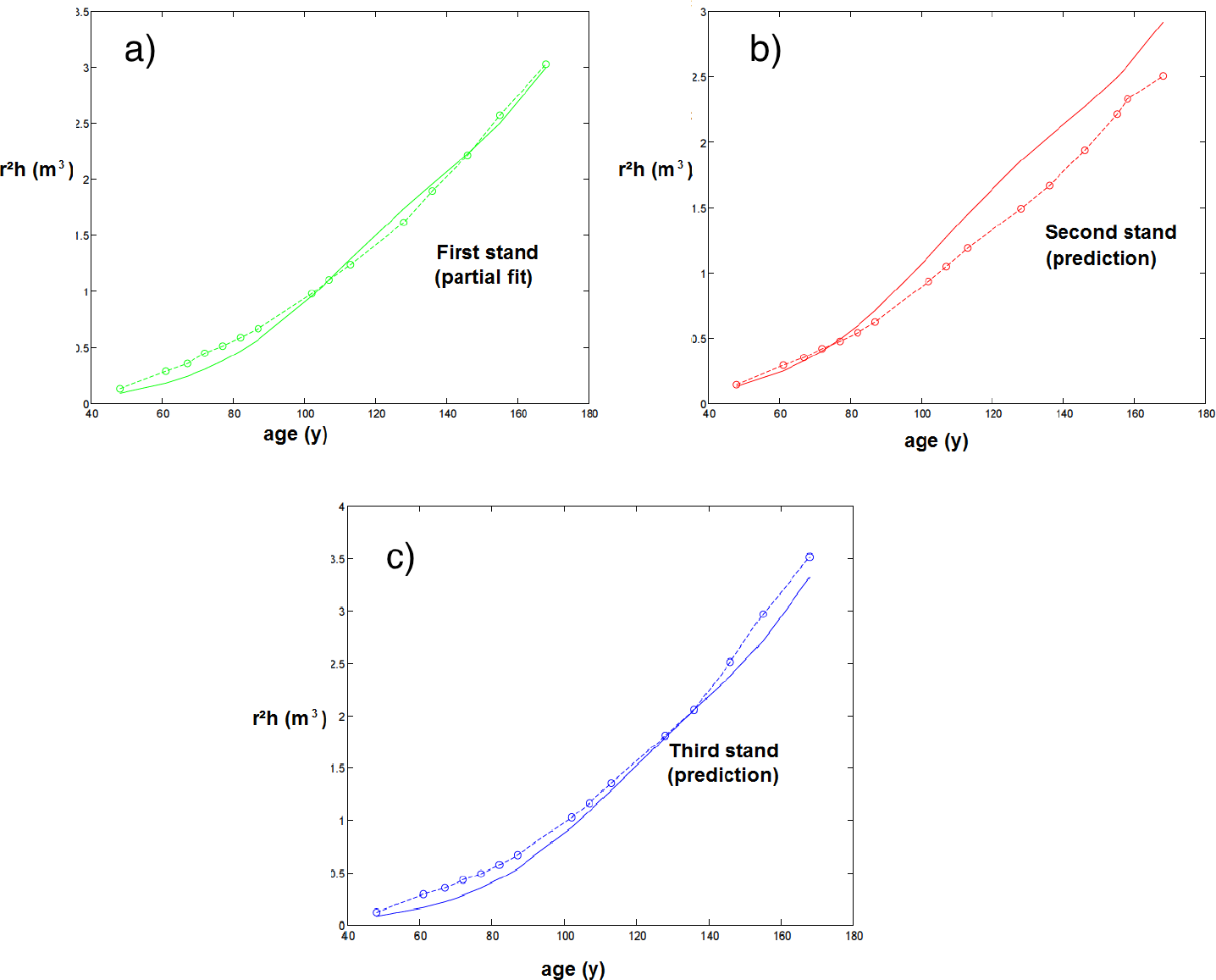
Plot of the stem volume with age at the first site. Continuous curves are computed with the model and start when the tree density is lower than 1,000 tree ha^-1^, while circles correspond to experimental measurements. a) and b) represent the results of the prediction with the parameters obtained by partial fitting. c) represents the partial fit on the reference stand.

**Figure 4.**
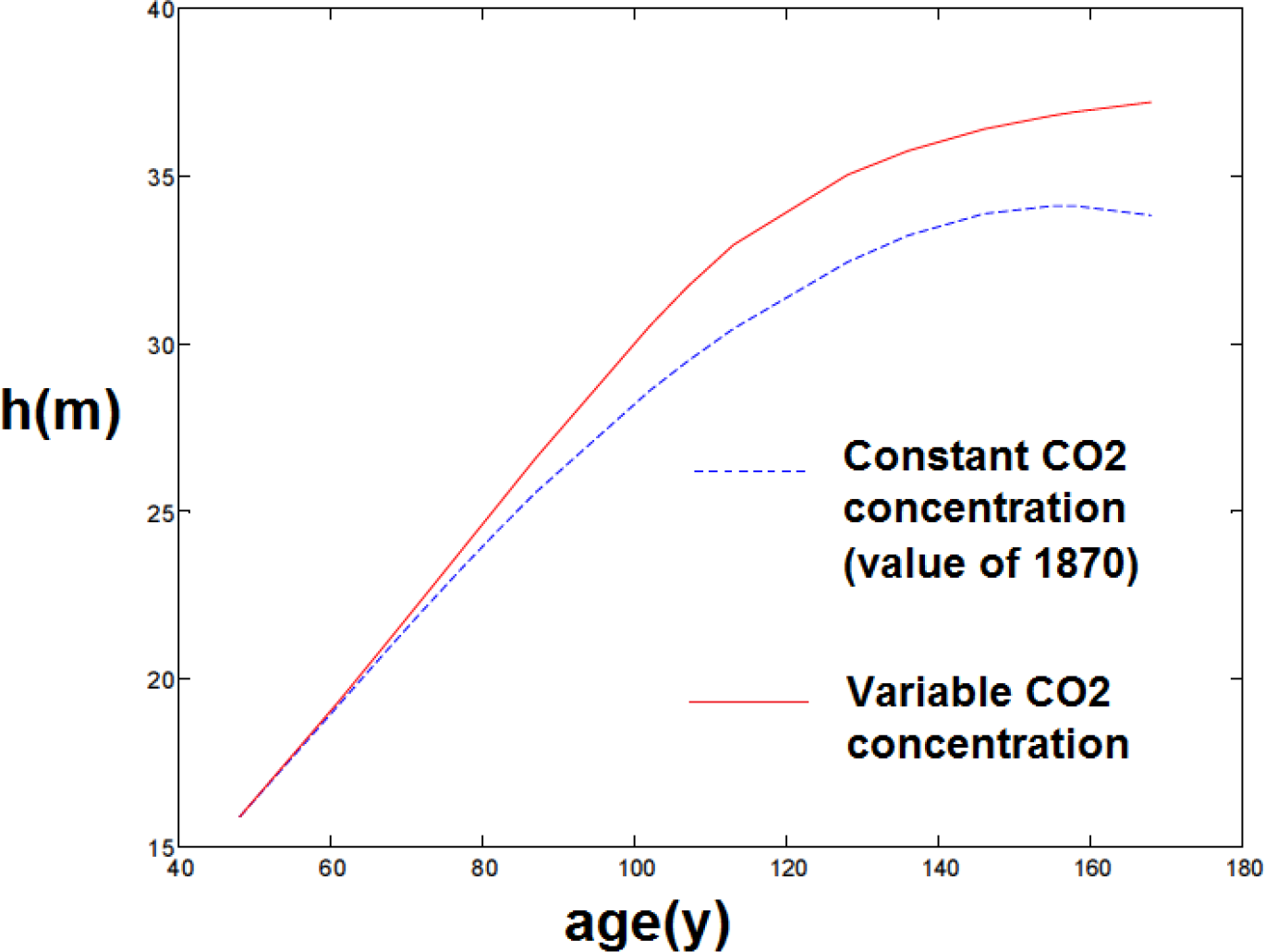
Comparison of simulations with and without increasing atmospheric CO_2_ and changing temperature from 1870.

## 4 Discussion

Our model starts from physiological considerations about trees and several hypotheses to derive differential equations that are solved over time to obtain the evolution of a tree and predict its size. Although the model is fairly simple and far from taking every possible parameter of the tree into account, it seems to obtain very good results and predictions which agree closely with observations from different, independent, stands of trees with different surrounding conditions.

Interestingly, our approach gives rise to two limiting tree heights: *h*_1_, the limit height above which photosynthesis cannot occur, and *h*_2_, the height above which growth is limited by the capacity of the meristematic regions themselves to grow at low water potential. These two heights represent different processes: *h*_1_ includes the factors that limit photosynthesis such as minimal turgor potential and possible xylem cavitation (i.e. source limitations), whereas *h*_2_ takes into account intrinsic meristem factors such as the physiological limits of cell division and growth limits such as cell wall extension when the water potential is too low (i.e. sink limitations).

While *h*_1_ (and, more generally, source processes) is often considered as the only limiting height to explain the maximal height of trees (e.g. [23]), under some external conditions of light and temperature it could be that *h*_2_ (i.e. sink activity) becomes limiting (cf. [22]). Although in this model, water potential is still the critical factor which limits the height of trees, the amount of light received and the temperature during the growth period also play roles that could produce one or other limit and influence its value. Hence, our approach could give a new and more complete understanding of the limits of tree height, and growth more generally, through a balanced consideration of both source and sink limitations.

As a first dynamic model for an individual tree using such considerations about control mechanisms and the maximal meristem-sustained limited rate of growth, improvements could probably be made by introducing additional parameters, especially taking the roots into account explicitly, and improving the function of the red:far-red ratio. Nevertheless, the model already gives very good results while being relatively simple, allowing its wider use such as through allowing many trees to interact with each other via shading, water potential, etc., and obtaining a model for the whole forest. So far the partial fitting of the parameters within bounds of 20% takes less than 120 s on a desk-top computer, and computing height over 100 yr less than 0.1 s, which suggests that its use as a growth simulator within a forest model could be achieved with today’s computer resources.

The relative simplicity of the model and its extremely encouraging results should stimulate further use of similar differential models in the future. Starting from physiological considerations to derive differential equations and an individual tree model as a cornerstone for simulating a whole forest may considerably improve our understanding of forest behaviour, including, but not limited to, the prediction of the overall reaction of forests to increasing CO_2_ concentrations in order to address the future terrestrial carbon sink.

To conclude, we have proposed a differential model for modelling tree growth over time under external conditions such as temperature, soil moisture, and CO_2_ concentration. This model takes into account not only photosynthesis and the carbon balance but also meristem behaviour and cellular growth limits. We established a procedure to parametrise the model with measurable quantities and reference measurements. This model seems not only to fit data very well but also to give accurate predictions for tree height and tree volume.

## 5. Acknowledgements

Amaury Hayat wishes to thank the Cambridge Faculty of Mathematics for support via the PMP scheme. All authors thank those involved in the measurements of the plot data at the two experimental sites.

## References

1. Bonan GB. Forests and Climate Change: Forcings, Feedbacks, and the Climate Benefits of Forests. Science. 2008 Jun;320(5882):1444–1449. Available from: http://www.sciencemag.org/content/320/5882/1444.abstract.

2. Pan Y, Birdsey RA, Fang J, Houghton R, Kauppi PE, Kurz WA, et al. A Large and Persistent Carbon Sink in the World’s Forests. Science. 2011 Aug;333(6045):988–993. Available from:http://www.sciencemag.org/cgi/doi/10.1126/science.1201609.

3. Cramer W, Bondeau A, Woodward FI, Prentice IC, Betts RA, Brovkin V, et al. Global response of terrestrial ecosystem structure and function to CO2 and climate change: results from six dynamic global vegetation models. Global Change Biology. 2001 Apr;7(4):357–373. Available from: http://dx.doi.org/10.1046/j.1365-2486.2001.00383.x.

4. Anav A, Friedlingstein P, Kidston M, Bopp L, Ciais P, Cox P, et al. Evaluating the Land and Ocean Components of the Global Carbon Cycle in the CMIP5 Earth System Models. Journal of Climate. 2013 Mar;26(18):6801–6843. Available from: http://dx.doi.org/10.1175/JCLI-D-12-00417.1.

5. Friend AD, Lucht W, Rademacher TT, Keribin R, Betts R, Cadule P, et al. Carbon residence time dominates uncertainty in terrestrial vegetation responses to future climate and atmospheric CO_2_. Proceedings of the National Academy of Sciences. 2014 Mar;111(9):3280–3285. Available from: http://www.pnas.org/lookup/doi/10.1073/pnas.1222477110.

6. Korner C. Carbon limitation in trees. Journal of Ecology. 2003 Feb;91(1):4–17. Available from: http://doi.wiley.com/10.1046/j.1365-2745.2003.00742.x.

7. Fatichi S, Leuzinger S, Korner C. Moving beyond photosynthesis: from carbon source to sink-driven vegetation modeling. New Phytologist. 2014 Mar;201(4):1086–1095. Available from: http://dx.doi.org/10.1111/nph.12614.

8. Leuzinger S, Manusch C, Bugmann H, Wolf A. A sink-limited growth model improves biomass estimation along boreal and alpine tree lines. Global Ecology and Biogeography. 2013 Aug;22(8):924–932. Available from: http://onlinelibrary.wiley.com/doi/10.1111/geb.12047/abstract.

9. Schiestl-Aalto P, Kulmala L, Makinen H, Nikinmaa E, Makela A. CASSIA-a dynamic model for predicting intra-annual sink demand and interannual growth variation in Scots pine. New Phytologist. 2015 Apr;206(2):647–659. Available from: http://doi.wiley.com/10.1111/nph.13275.

10. Gea-Izquierdo G, Guibal F, Joffre R, Ourcival JM, Simioni G, Guiot J. Modelling the climatic drivers determining photosynthesis and carbon allocation in evergreen Mediterranean forests using multiproxy long time series. Biogeosciences. 2015 Jun;12(12):3695–3712. Available from: http://www.biogeosciences.net/12/3695/2015/.

11. Aloni R. Differentiation of Vascular Tissues. Annual Review of Plant Physiology. 1987 Jun;38(1):179–204. Available from: http://www.annualreviews.org/doi/ abs/10.1146/annurev.pp.38.060187.001143

12. Ritchie GA. Evidence for red:far red signaling and photomorphogenic growth response in Douglas-fir (Pseudotsuga menziesii) seedlings. Tree Physiology. 1997 Mar;17(3):161–168. Available from: http://treephys.oxfordjournals.org/content/17/3/161.abstract

13. Brackmann K, Greb T. Long-and short-distance signaling in the regulation of lateral plant growth. Physiologia Plantarum. 2014 Jun;151(2):134–141. Available from: http://doi.wiley.com/10.1111/ppl.12103

14. Aloni R. Ecophysiological implications of vascular differentiation and plant evolution. Trees. 2015 Feb;29(1):1–16. Available from: http://dx.doi.org/10.1007/s00468-014-1070-6

15. Franklin KA. Shade avoidance. New Phytologist. 2008 Sep;179(4):930–944. Available from: http://dx.doi.org/10.1111/j.1469-8137.2008.02507.x

16. Morgan DC, Smith H. Linear relationship between phytochrome photoequilibrium and growth in plants under simulated natural radiation. Nature. 1976 Jul;262(5565):210–212. Available from: http://dx.doi.org/10.1038/262210a0

17. Aphalo PJ, Ballare CL, Scopel AL. Plant-plant signalling, the shade-avoidance response and competition. Journal of Experimental Botany. 1999 Nov;50(340):1629–1634. Available from: http://jxb.oxfordjournals.org/content/50/340/1629.abstract

18. Li J, Li G, Wang H, Wang Deng X. Phytochrome Signaling Mechanisms. The Arabidopsis Book / American Society of Plant Biologists. 2011 Aug;9. Available from: http://www.ncbi.nlm.nih.gov/pmc/articles/PMC3268501/

19. Precht H,Christophersen J, Hensel H, Larcher W. Temperature and Life. New York, Heidelberg, and Berlin: Springer-Verlag; 1973.

20. Deleuze C, Houllier F. A Simple Process-based Xylem Growth Model for Describing Wood Microdensitometric Profiles. Journal of Theoretical Biology. 1998 Jul;193(1):99–113. Available from: http://linkinghub.elsevier.com/retrieve/pii/S0022519398906890

21. Campioli M, Gielen B, Gockede M, Papale D, Bouriaud O, Granier A. Temporal variability of the NPP-GPP ratio at seasonal and interannual time scales in a temperate beech forest. Biogeosciences. 2011;8(9):2481–2492. Bibtex: bg-8-2481-2011. Available from: http://www.biogeosciences.net/8Z2481/2011/

22. Friend AD. The prediction and physiological significance of tree height. In: Vegetation Dynamics and Global Change; 1993. p. 101–115.

23. Koch GW, Sillett SC, Jennings GM, Davis SD. The limits to tree height. Nature. 2004 Apr;428(6985):851–854. Available from: http://dx.doi.org/10.1038/nature02417

24. Du N, Fan J, Chen S, Liu Y. A hydraulic-photosynthetic model based on extended HLH and its application to Coast redwood (Sequoia sempervirens). Journal of Theoretical Biology. 2008 Jul;253(2):393–400. Available from: http://linkinghub.elsevier.com/retrieve/pii/S002251930800129X.

25. Dormand JR, Prince PJ. A family of embedded Runge-Kutta formulae. Journal of Computational and Applied Mathematics. 1980 Mar;6(1):19–26. Available from: http://linkinghub.elsevier.com/retrieve/pii/0771050X80900133.

26. Zianis D, Mencuccini M. Aboveground net primary productivity of a beech (Fagus moesiaca) forest: a case study of Naousa forest, northern Greece. Tree Physiology. 2005 Jun;25(6):713–722. Available from:http://treephys.oxfordjournals.org/content/25/6Z713.abstract.

23. Holzwarth F, Kahl A, Bauhus J, Wirth C. Many ways to die-partitioning tree mortality dynamics in a near-natural mixed deciduous forest. Journal of Ecology. 2013 Jan;101(1):220–230. Available from:http://doi.wiley.com/10.1111/1365-2745.12015

23. Pretzsch H, Biber P, Schutze G, Uhl E, Rotzer T. Forest stand growth dynamics in Central Europe have accelerated since 1870. Nature Communications. 2014 Sep;5:4967. Available from:http://www.nature.com/doifinder/10.1038/ncomms5967

23. Nagel TA, Levanic T, Diaci J. A dendroecological reconstruction of disturbance in an old-growth Fagus-Abies forest in Slovenia. Annals of Forest Science. 2007Jan;64(8):891–897. Available from: http://link.springer.com/10.1051/forest:2007067

